# Cascading indirect genetic effects in a clonal vertebrate

**DOI:** 10.1101/2021.02.27.433187

**Authors:** Amber M. Makowicz, David Bierbach, Christian Richardson, Kimberly A. Hughes

## Abstract

Understanding how individual differences among organisms arise and how their effects propagate through groups of interacting individuals are fundamental questions in biology.Individual differences can arise from genetically-based variation in the conspecifics with which an individual interacts, and these effects might then be propagated to other individuals. Using a clonal species, the Amazon molly (*Poecilia formosa*), we test the hypothesis that such indirect genetic effects (IGE) propagate beyond individuals that experience them firsthand. We tested this hypothesis by exposing genetically identical Amazon mollies to social partners of different genotypes, and then moving these individuals to new social groups in which they were the only member to have experienced the IGE. We found that genetically different social environments induced different levels of aggression experienced by the focal animals, and that these genetically-based social effects carried over into new social groups to influence the behavior of individuals that did not directly experience the previous social environments. Our data reveal that IGE can cascade beyond the individuals that directly experience them to influence phenotypes even when there is no genetically-based variation present within interacting groups. Theoretical and empirical expansion of the quantitative genetic framework developed for IGE to include cascading and other types of carry-over effects will improve understanding of social behavior and its evolution.

## 1. Introduction

The environment an individual experiences includes its interactions with conspecifics, and individual differences have long been known to influence these interactions [1, 2]. Understanding how these individual differences arise and how their effects propagate through groups are fundamental questions in biology [2-4]. One cause of both individual variation and propagation of effects in groups are indirect genetic effects (IGE) [4-7]. IGE arise when individual phenotypes are influenced by genetically-based phenotypic differences in conspecific partners, and they have been documented to affect behavioral, life history, and morphological traits in a wide variety of taxa [e.g., 3, 8-17]. For example, behavior and body condition of mosquitofish is influenced by genetically-based differences in body color of their social partners [18, 19], and behavioral, physiological, and morphological traits in laboratory mice are influenced by genotypes of their social partners [20]. Much of the empirical IGE literature focuses on how partner genetic variation influences phenotypes of focal individuals. While understanding these dyadic interactions is important, much less is known about IGE on group-level characteristics or the degree to which IGE can propagate to affect phenotypes of individuals that do not experience them firsthand. Because IGE can profoundly affect phenotypes, fitness, and the rate and direction of evolution [21-23], understanding possible cascading or carry-over effects within groups is necessary to understand phenotypic variation and evolution.

There have been two studies, to our knowledge, that have investigated IGE beyond those caused by dyadic interactions. The first used fruit flies (*Drosophila melanogaster*), to measure first-order IGE on male aggressive behavior (i.e., how the genotype of stimulus individuals influences the phenotypes of individuals with which they interact) and second-order IGE (effects of the stimulus genotypes on the interaction between two other members of the group) [24]. In that study, Saltz showed that differences in aggressive behavior between two stimulus genotypes had first-order effects on aggressive behavior in their partners and had second-order effects on aggressive interactions between other group members. The second study, also using

*D. melanogaster*, reported that the genotype of stimulus individuals influenced emergent, group-level behavior of focal individuals [25]. Specifically, Anderson et al. reported that, if individuals of the stimulus genotype were closer to one another, on average, then the focal individuals were also more cohesive, and interactions between stimulus and focal individuals were less frequent.

Together, these two experiments indicate that IGE can extend beyond the direct effects of one individual on another. However, it remains unknown whether IGE previously experienced by one or a few group members can influence phenotypes of individuals that were never exposed to the IGE. That is, can IGE “cascade” beyond individuals that experience them firsthand? Previous work in animals indicates that individual group members can influence group behavior [1, 26-29]. However, this literature has generally not focused on prior social experience as a factor that generates differences between influential group members [but see 30-32]; moreover, we know of no studies that implicate IGE as a cause of such differences. It is challenging to measure prior influence of IGE because it is difficult to replicate group genotypic composition and genetically-based differences in social environment in sexually-reproducing species. Nevertheless, many organisms exhibit either dispersal or fission-fusion social structure, so understanding IGE caused by prior social environments is critical to understanding the evolution of social behaviors.

Naturally clonally-reproducing organisms provide an opportunity to measure these effects outside of model species and without inbreeding or complex breeding designs. The Amazon molly (*Poecilia formosa*) is a gynogenetic, all-female species [33] that arose from a single hybridization event between a male sailfin molly (*Poecilia latipinna*) and a female Atlantic molly (*Poecilia mexicana*) about 100,000 generations ago [34, 35]. Although reproduction is clonal, females require sperm from a male of one of the ancestral species (sailfin or Atlantic molly) to initiate embryogenesis of unreduced ova [36]. Many distinct clonal lineages arose from the original diploid lineage through mutation or complete and/or partial incorporation of paternal genetic material, which can be stable and transmitted to subsequent generations [36, 37]. This accumulation of genetic diversity in a gynogenetic species produces groups in which social interactions occur on multiple levels: within-clone interactions, among-clone interactions, and interspecies interactions between Amazons and their sexual hosts. While the interactions between Amazons and their hosts have been the focus of many investigations over the past forty years [e.g., 33, 38-41], little attention has focused on the social interactions within [42] and among the different clonal lineages; however, previous research does suggest that clonal lineages vary in the social behaviors [43].

In natural populations, the number of clonal lineages that co-occur can vary dramatically from a single lineage to more than a dozen [44-46]. Therefore, the degree of competition and the frequency with which females encounter conspecifics of different lineages can vary greatly across time and space. One of the first studies to investigate social behaviors among different clones reported that females could distinguish between lineages, associate preferentially with fish of their own lineage, and were more aggressive toward unrelated clones [43]. Other studies have reported that different features of the social environment can influence social behavior, especially aggression, within and among clonal lineages, including early dominance interactions [47] and the degree of familiarity among individuals [42, 48]. These data suggest that individual behavior depends in part on the clonal composition of the social environment; that is, IGE likely regulate phenotypic variation and social dynamics in natural populations.

We used clonal variation in Amazon mollies to test the hypothesis that IGE propagate beyond individuals that experience them firsthand. This hypothesis predicts that variation in behavior generated by IGE in a previous social environment will influence the behavior of naïve individuals when an animal with this prior experience joins their group. To distinguish this effect from first- and second-order IGE, we use the term ‘cascading IGE’. Based on extensive literature indicating that individual differences in behavior affects group-emergent phenotypes [reviewed by 1], we also predicted that cascading IGE will influence group-emergent behavior in these fish. We tested these predictions by exposing genetically identical Amazon mollies to social partners of different genotypes and then moving these individuals to new social groups in which they were the only member to have experienced IGE. This experimental paradigm simulates the fission-fusion dynamics often observed in poeciliid fishes in the natural environment [49-51].

## 2. Material and methods

### (a) Study Specimens

Three distinct clonal lineages were used in this study, each descended from individuals collected from the Río Purificacíon in Nuevo Padilla, Mexico (24°4′42.85′′N, 99°7′21.76′′W), and maintained in a greenhouse at the Mission Road Research Facility of Florida State University. Both Clone 1 (Schartl) and Clone 2 (AMM#11) are diploid with microchromosomes, although the microchromosomes are distinctly different between the two lineages [43, 52]. The focal clone (3N) is a triploid without any microchromosomes, this clonal lineage was chosen at random to be the focal clone. Details concerning fish care can be found in the electronic supplementary materials, Methods.

### (b) Long-term social environments

Focal females were placed into 18.9L aquaria in one of three different long-term social environments: (1) 1 focal female + 2 sister clones; (2) 1 focal female + 2 females from a Clone 1; and (3) 1 focal female + 2 females from Clone 2. That is, each aquarium contained 1 focal fish and 2 “social partner” fish. Females placed into the Monoclonal social environment were unfamiliar with each other as they originated from different rearing and recovery tanks prior to the start of the experiment. The partner fish genotypes, but not the genotype of focal fish, differed among treatments. Each social-environment treatment was replicated 12 times for a total of 36 experimental tanks. Experimental tanks were set up using a randomized complete block design (one replicate of each treatment per block) over the course of two weeks (6 blocks set up per week) until all 12 blocks were complete. All females ranged between 27 and 38 mm in body length with a maximum size difference among females within each social environment of 4 mm to reduce the influence of body size on aggression [48].

To characterize differences in the social environment induced by the three different social treatments, we measured social interactions in the experimental tanks at 9 different times over the course of the experiment: 10 min after placing the focal fish in the social environment (week 0), weekly for the first four weeks thereafter (weeks 1-4), and then biweekly until a total of 12 weeks of exposure (weeks 6, 8, 10, and 12). Behavior measured at week 0 represents a baseline because females had no prior exposure to experimental social environments at this time point. Social behavior in the experimental tanks consisted mainly of aggressive interactions (bites, tail beats, and chasing); few affiliative or neutral behaviors (e.g., swimming in the same direction or foraging simultaneously within 2 body lengths) were observed outside an aggressive context (e.g., proceeding or following biting, chasing or tail beating). We counted the number of bites and tail beats performed and the total time spent performing these behaviors and chasing other females. Tail beats were rarer than bites, and the distribution was zero-inflated. We therefore summed the total number of bites and tail beats observed, and separately summed the total time spent in these aggressive interactions to produce two overall measures of aggression: total number of aggressive acts and total time spent in aggression. Both measures were log-transformed before analysis, after adding 1 to account for zero values. These assays were recorded by a live observer blind to the treatments for a duration of 10 minutes.

### (c) Naïve-group tests

Each focal female was introduced to a pair of novel (‘naïve’) social partners three times over the course of the experiment (at 0, 4, and 12 weeks). A different pair of naïve social partners was used at each of these trials, and those partner fish were not used with any other focal female. We measured the average behavior of these naïve-groups before exposing focal fish to genetically different long-term social environments (week 0) and after 4- and 12-weeks of exposure (see Figure 1). To do so, individual focal females were removed from their rearing tank (at week 0) or their long-term social environment tank (at weeks 4 and 12) and placed in a “naïve-group” test chamber with two unfamiliar females from the same clonal lineage as the focal fish, size matched to the focal fish (± 4mm), and in the same reproductive state. These novel fish were drawn from monoclonal, non-breeding rearing tanks similar to those from which focal and stimulus females originated and were, therefore, not exposed to the experimental social environments experienced by the focal females. After we introduced the focal fish into the naïve-group test chamber, we video recorded all three fish for 10 minutes, after which the focal female was removed and placed back into her experimental social environment (Figure 1).

**Figure 1:**
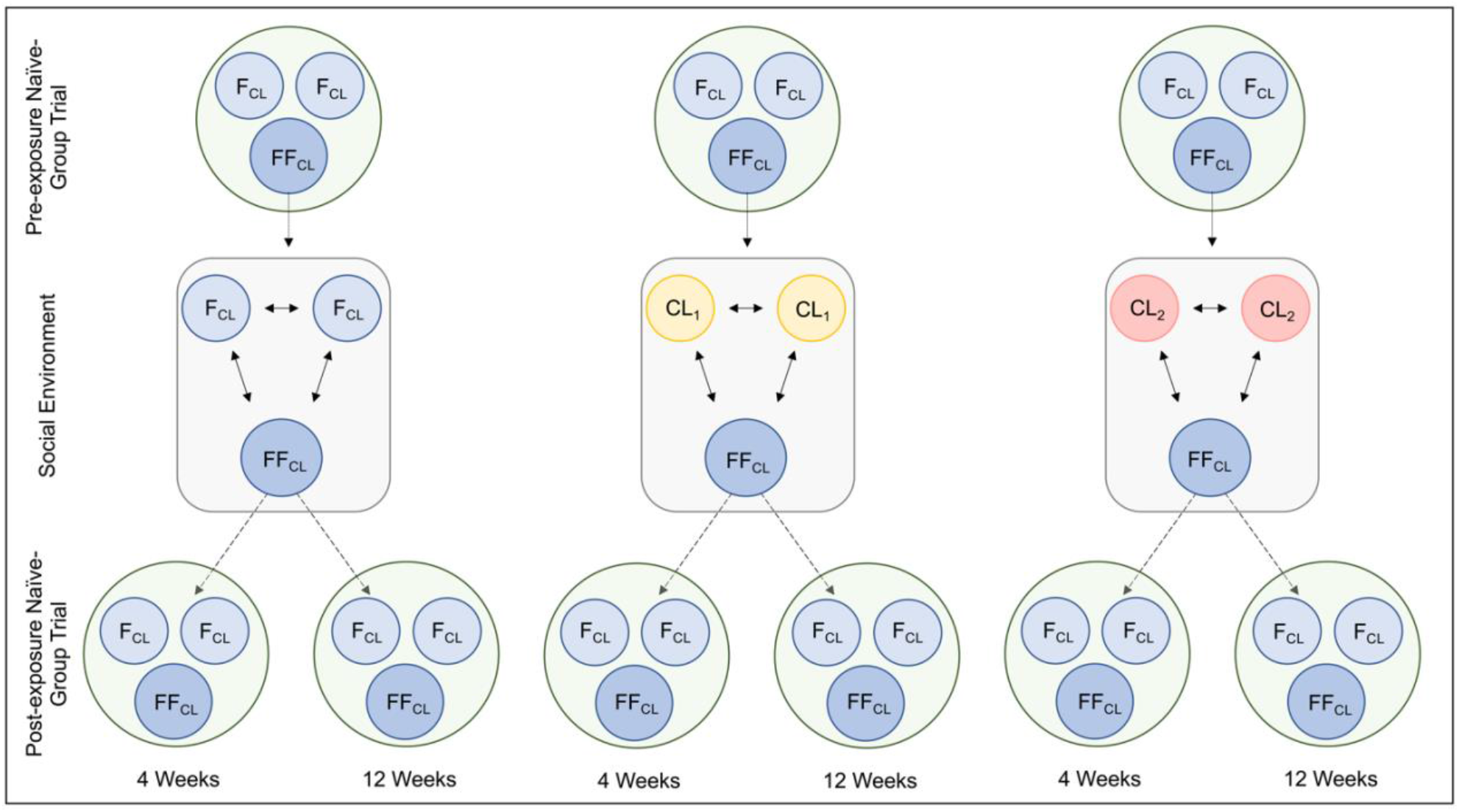
Schematic of experimental design illustrating the focal females (FF_CL_) tested for pre-exposure exploratory behaviors with two novel sister clones (F_CL_) at week 0. Focal females were then transferred into one of the three different social environments: Monoclonal (FF_CL_ + 2 F_CL_); Clone 1 (FF_CL_ + 2 CL_1_); or Clone 2 (FF_CL_ + 2 CL_2_). After 4 weeks and then again at 12 weeks of exposure to these social environments, the exploratory behaviors of the focal females were tested again with novel F_CL_ individuals. Note that the F_CL_ partners of the FF_CL_ were different individuals at each time period. That is, each individual F_CL_ was included in only one trial.

Because we were interested in emergent (group-level) behavior of the naïve groups, the naïve-group test chamber was an open field, circular tank (55.9 cm diameter), with half the bottom and corresponding sides painted white and the other half grey. In the center of the frame, a camera (JVC Everio 1920×1080 HD video camcorder) was suspended 1.1 m above the tank. All videos were 6 minutes long and were analyzed by a blind observer using EthoVision XT (Noldus, v14). For more information regarding video recording, editing, and EthoVision, see the electronic supplementary materials, Methods.

Although fish could be individually tracked, the focal individual could not be distinguished from the novel partner fish on the videos; therefore, we did not calculate separate metrics for focal and novel partner fish. We interpret behaviors as reflecting stress-related behavior or tendency to be exploratory. More stressed individuals are less active, travel shorter distances at lower velocity, spend more time frozen and in the grey zone (negative phototaxis), and are closer together; less stressed individuals tend to be more exploratory and cover more distance, move at higher velocity, enter zones more frequently, spend more time in the white zone and less time frozen, and have more distance between individuals [53, 54]. We also gathered baseline data on these behaviors by following the same procedure at the start of the experiment, before the focal fish had experienced the experimental social treatments (week 0; Figure 1: Pre-exposure).

### (d) Ethics

This research was approved by the Institutional Animal Care and Use Committee of Florida State University (1704 and 201900038).

### (e) Analyses

There were no significant differences in size (SL) among focal females in different social treatment groups, nor treatment-associated differences in size among the social partner fish used in the long-term and naïve-group trials (electronic supplemental material, Table S1). Nevertheless, we included SL of focal and partner fish as covariates in subsequent analyses because there was a non-significant trend for Clone 1 and Clone 2 social partner fish to differ in SL (electronic supplemental material, Table S2).

#### (e.1) Long-term social environment groups

We assessed the correlation structure of the two measurements of aggression to determine if they could be adequately represented by principal components (PC), and then used the first PC from this analysis as our measure of aggression (see Results). To determine if aggression was influenced by social treatment group, we used this PC1 score as the dependent variable in linear mixed models. In addition to the social treatment group, initial models included fixed effects of exposure time (weeks), treatment-by-time interaction, the baseline (week 0) measure of aggression PC1, and focal female standard length (log-transformed. A random group ID effect was used to account for repeated measures on groups (at weeks 4 and 12). Except where noted, this and other analyses of linear mixed models were conducted using SAS *Proc Glimmix* in SAS v. 9.4 [55] with a Gaussian error distribution and an identity link function. Post hoc comparisons of treatment group means were conducted using the simulation method of [56], as implemented by using the *adjust=simulate* option. See the electronic supplementary materials, Methods for more details on statistical models.

#### (e.2) Naïve-group tests

To determine the extent to which the presence of the focal individual influenced behavior in the naïve-groups, and thus to measure cascading IGE, we calculated two kinds of metrics: those that described average behavior of the 3 members of the group, and those that described individual behavior of fish within the group. For both analyses, we summarized six of the movement variables (distance traveled (cm), velocity (cm/s^2^), frequency entering white zone (count), duration in white zone (s), latency to enter white zone (s), time spent immobile (s; freezing behavior) with PC scores (see below). Distance between individuals (cm; shoaling distance) was analyzed separately (see below).

##### Average behavior of naïve-groups

We first assessed the correlation structure of the 7 behaviors to determine if they could be adequately represented by principal components. The six behaviors that described movement or physical position in the enclosure were all moderately to highly correlated with one another (0.4 < |r| < 1.0), but they were not correlated with the average shoaling distance between fish (all |r| < 0.2) (electronic supplementary material, Figure S1A), indicating that a PCA should include the 6 movement/position variables, but that shoaling distance should be analyzed separately. We used the first PC from this analysis as our measure of the movement and position of fish (see Results), and we used the log-transformed average shoaling distance as a measure of a group cohesion, since it arises from the relative positions of all three members of the group.

To determine if these two measures of naïve-group behavior were differentially affected by the social environment experienced by a single member of the group, we used the values at weeks 4 and 12 as the dependent variable in linear mixed models. Neither the size-related fixed effects nor the measures of aggression approached significance in the initial models (electronic supplemental material, Table S3B and C, Methods), so only treatment and exposure time (and their interaction) were retained in the final models. A random effect with focal female ID as the subject was used to account for the different naïve-group trials in which each focal female was used. Post hoc comparisons of group means were conducted as described above. We calculated the proportion of variance explained by the long-term social environment using the method of Jaeger et al. [57].

Since treatment groups varied in aggression (see Results), any overall association between aggression experienced in the long-term social environment and the behavior of naïve groups could have been obscured by the treatment effect in the models described above. To assess the overall relationship between aggression in the long-term environment and naïve group behavior, we therefore fit models for exploratory PC1 and shoaling identical to those described above, but with the only predictor variable being cumulative aggression in the long-term environment.

Focal females were tested in naïve groups once at baseline and twice more after 4 and 12 weeks of exposure to their long-term social groups. To assess the consistency of behavior of the naïve groups that contained the same focal female after 4 and 12 weeks of experience in their long-term social environments, we calculated the Pearson’s correlation between PC1 scores (or shoaling) at the two time periods [58]; we calculated 95% confidence limits using the z-transformation method.

##### Behavior of individuals in naïve-groups

The main purpose of this analysis was to determine if differences in the average behavior among naïve groups was attributable to all members of a group behaving similarly or to specific individuals within the group. For example, if the behavior of the three females within a group was very similar, then average differences among groups reflect the behavior of all group members. Alternately, if individuals within groups behaved differently from one another, then between-group differences could have been driven by the divergent behavior of a single group member. The former, but not the latter, would support cascading IGE because it would indicate that non-focal behavior was influenced by the prior social experience of the focal fish. Our primary measure of similarity of the behavior of individuals within naïve-groups was the ICC.

We first investigated the correlation structure of the same 8 behaviors described above but measured on individuals rather than the mean of the 3 fish in a group. As in the group-average data, the position/movement variables were moderately to highly correlated with each other, but not with shoaling distance (electronic supplementary material, Figure S1B). We therefore summarized the movement/position behavior of individual fish using the first PC of the 6 movement/position metrics (Table 1). We then calculated the ICC of the individual exploratory behavior scores using a linear mixed model with a random effect corresponding to group ID. The ICC was estimated as the ratio of among-group variance to the total variance and confidence intervals were determined using parametric bootstrap estimates [58].

**Table 1.** PC loadings for PC1 on group-averaged and individual-level exploratory/stress behaviors.

| Level | Model | Measurement | PC1 loading |
| --- | --- | --- | --- |
| Group-averaged | PC1 exploratory/stress | Total distance traveled | 0.951 |
|  |  | Velocity | 0.951 |
|  |  | Frequency entering white zone | 0.956 |
|  |  | Duration in white zone | 0.686 |
|  |  | Latency to enter white zone | -0.739 |
|  |  | Time spent frozen in place | -0.899 |
| Individual-level | PC1 exploratory/stress | Total distance traveled | 0.949 |
|  |  | Velocity | 0.948 |
|  |  | Frequency entering white zone | 0.937 |
|  |  | Duration in white zone | 0.642 |
|  |  | Latency to enter white zone | -0.702 |
|  |  | Time spent frozen in place | -0.897 |

## 3. Results

### (a) Summary measures of aggression and movement explain most of the variation

The two measures of aggression (number of acts and time spent) were highly correlated (R^2^=0.803, p<0.0001), with the first PC explaining 96.9% of the total variation. For the average behaviors of the naïve-groups, the first PC summarizing the 6 movement/position variables explained 75.79% of the total variation, and it was the only PC with an eigenvalue >1 (Table 1). Behaviors associated with exploration loaded positively on PC1 (distance, velocity, and duration in the white zone, and frequency entering white zones), while behaviors associated with stress loaded negatively on PC1 (freezing, latency to enter white zone, Table 1). We therefore considered positive values of PC1 to indicate a tendency to explore, and negative values to indicate lack of exploration or stress-like behaviors.

### (b) Long-term social environments differ in aggressive behavior

Long-term social groups in which the focal fish was housed with two females of her own clonal lineage exhibited more aggression than groups where the social partners were Clone 1 or Clone 2 fish (Figure 2A, Table 2A, effect estimates provided in electronic supplementary material, Figure S2 and Table S4). On average, fish in the Monoclonal environment performed 60% more aggressive acts than fish in the Clone 1 environment (14.45±1.49 vs. 9.04±1.27 acts per 10-minute observation bout, respectively; fish in the Clone 2 environment performed 11.04±1.18 aggressive acts per bout, on average). Post hoc tests indicated that the Monoclonal social environment elicited significantly more aggression than the Clone 1 environment (t_256.1_=3.93, p<0.001), but no other contrasts were significant after adjustment for multiple tests (Monoclonal vs Clone 2: t_248.8_=2.15, p=0.087; Clonal 1 vs Clone 2: t_247.1_=-2.04, p=0.114).

**Figure 2:**
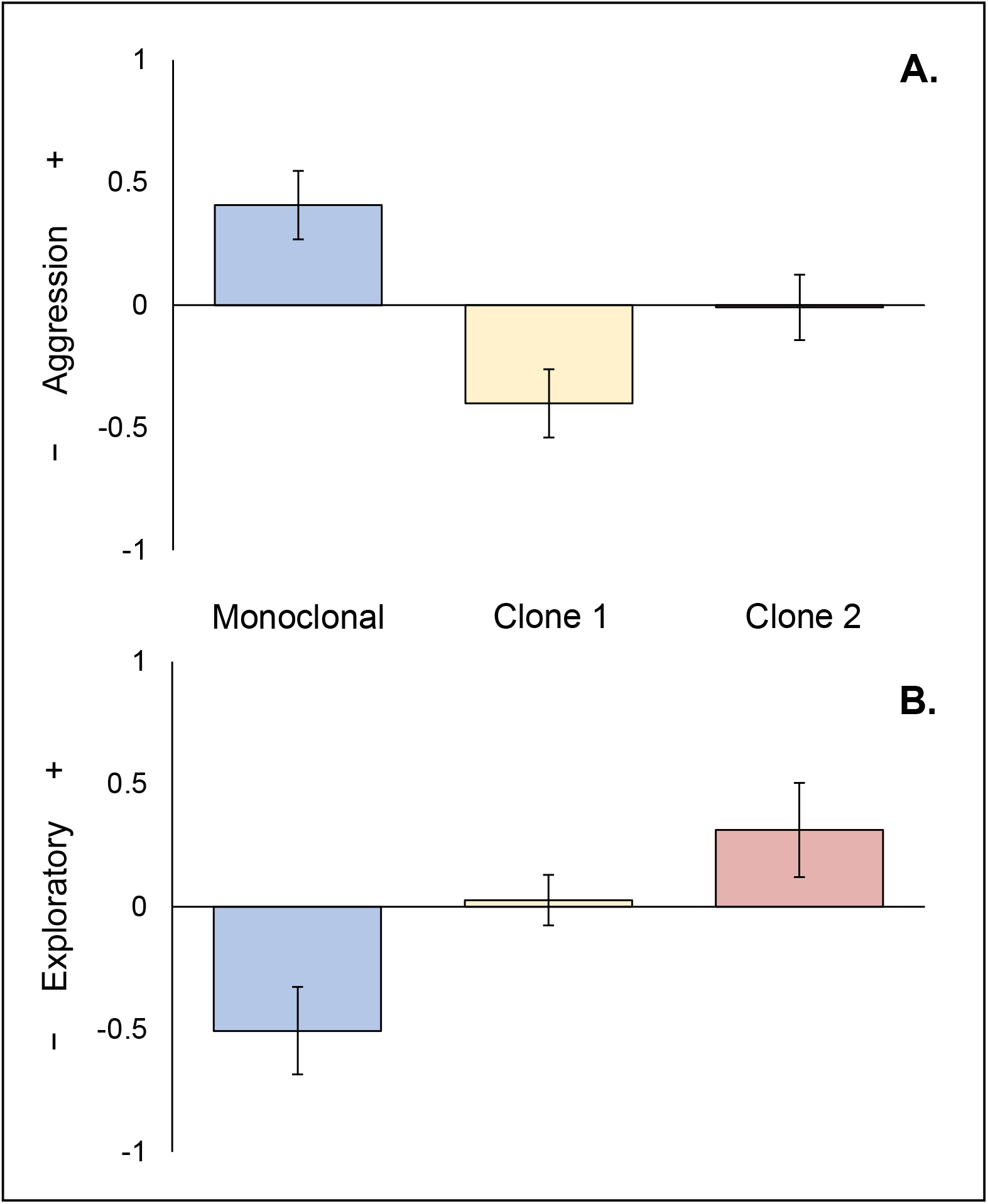
Least square means +/- standard error for (**A**.) aggression PC1 for each long-term social environment (Monoclonal (blue); Clone 1 (yellow); Clone 2 (pink)). Positive values indicate more aggression. (**B**.) PC1 for exploratory/stress behaviors in the naïve-group trials. Group-averaged exploratory behaviors with positive values indicating more exploratory behaviors and negative values indicate less exploratory and more stress behaviors.

**Table 2:** Proportion of variance explained, test statistic and p-value for statistical models of aggression PC1 (**A**.), exploratory/stress behaviors (PC1) (**B**.), shoaling behaviors (**C**.), the effects of cumulative aggression on exploratory/stress behaviors (PC1) (**D**.), and the effects of cumulative aggression on shoaling behaviors (**E**.).

| Model | Effect | Proportion of variance explained | Statistic | P-value |
| --- | --- | --- | --- | --- |
| *A. Aggressive behavior in long-term social treatments (PC1)* |  |  |  |  |
| | Focal female standard length | $R^2 = 0.015$ | $F_{1,208.1} = 3.26$ | 0.072 |
| | **Social environment** | $R^2 = 0.058$ | **$F_{2,250.3} = 7.70$** | **<0.001** |
| | Exposure time | $R^2 = 0.039$ | $F_{7,221.2} = 1.30$ | 0.253 |
| | Social environment*Time | $R^2 = 0.040$ | $F_{14,232.6} = 0.69$ | 0.788 |
| *B. Exploratory/stress behavior in naïve-group trials (PC1)* |  |  |  |  |
| | **Social environment** | $R^2 = 0.431$ | **$F_{2,19.46} = 5.13$** | **0.016** |
| | Exposure time | $R^2 = 0.023$ | $F_{1,19.4} = 1.11$ | 0.306 |
| | Social environment*Time | $R^2 = 0.089$ | $F_{2,18.13} = 0.58$ | 0.568 |
| *C. Shoaling distance in naïve-group trials* |  |  |  |  |
| | Social environment | $R^2 = 0.056$ | $F_{2,20.5} = 0.25$ | 0.779 |
| | Exposure time | $R^2 = 0.003$ | $F_{1,28.63} = 0.06$ | 0.816 |
| | Social environment*Time | $R^2 = 0.242$ | $F_{2,20.77} = 3.08$ | 0.067 |
| *D. Cumulative aggression on exploratory/stress behaviors in naïve-group trials (PC1)* |  |  | **$F_{1,19.66} = 5.42$** | **0.031** |
| *E. Cumulative aggression on shoaling distance in naïve-group trials* | | | $F_{1,26.3} = 1.19$ | 0.285 |

### (c) Genetic differences in prior social experience for one group member affected average behavior of the group

The long-term social environment experienced by a single focal fish affected the average exploratory behavior of the group when paired with otherwise naïve individuals (Table 2B, Figure 2B, and electronic supplementary material, Figure S3, effect estimates in electronic supplementary material, Table S5). Indeed, the social environment of the focal fish explained 43.1% of the total variation in the exploratory/stress PC1 scores. Groups in which the focal individual experienced the Monoclonal long-term social environment exhibited more stress-related behavior (negative values on exploratory PC1) than groups in which the focal individual experienced Clone 1 or Clone 2 social environments (post-hoc tests: Monoclonal vs Clone 1, t_17.58_=-2.59, p=0.044; Monoclonal vs Clone 2, t_21.88_=-3.12, p=0.015). Naïve-groups in which the focal fish had experienced social environments containing Clone 1 and Clone 2 did not differ significantly from each other after correction for multiple tests (t_16.81_=-1.31, p=0.405). This result is particularly striking because all focal and stimulus fish in this assay were members of a single clonal linage and, therefore, genetically identical. Surprisingly, the long-term social environment of focal animals not only affected the average behavior of naïve groups, it also affected the variance in behavior (i.e., the variance structure differed significantly between treatments; electronic supplementary material, Table S6).

Cumulative aggression was linearly related to exploratory/stress behaviors of the naïve-group trials in a model in which aggression was the only predictor (Table 2D). The more aggression a female experienced in her long-term social environment, the less likely she was to exhibit positive, exploratory behaviors and more likely to exhibit stress-like behaviors (β=-0.401±0.172). This result suggests that aggression might be a phenotype that influenced focal females and thus, the exploratory stress behaviors exhibited in the naïve-group trials.

The long-term social environment also affected the consistency in behavior over time of the naïve groups were (electronic supplementary material, Table S7). Specifically, the naïve groups with focal females originating from the Monoclonal social environment showed high consistency of exploratory behavior across weeks 4 and 12 (r=0.732; figure 3A), whereas naïve groups with focal females from the Clone 2 social environment exhibited much lower consistency (r=0.280), and groups with focal females from the Clone 1 long-term social environment exhibited a negative correlation between behaviors across time periods (r=-0.759).

**Figure 3:**
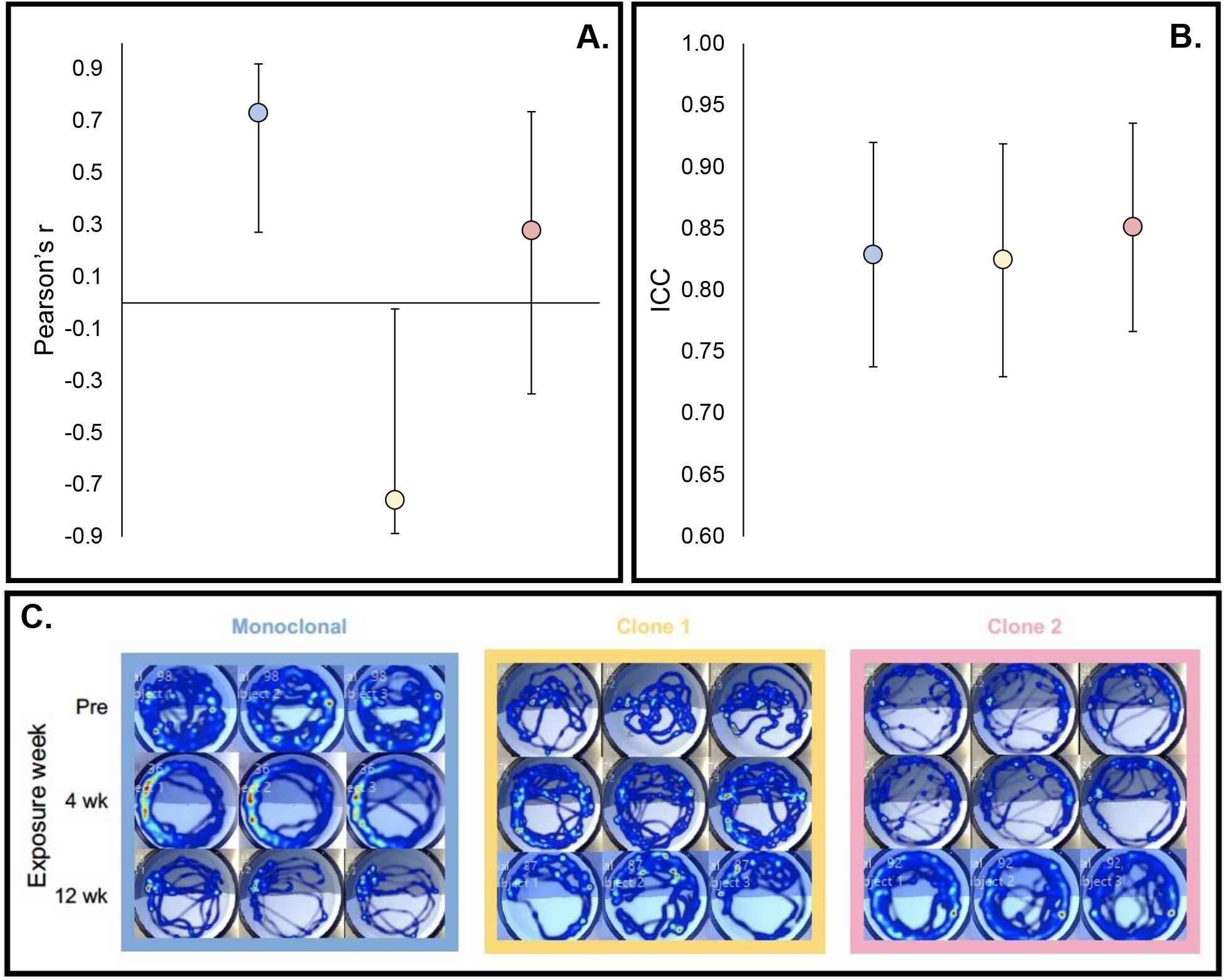
In each panel, the different colors represent Monoclonal (blue), Clone 1 (yellow), Clone 2 (pink). (**A**.) Consistency of behavior of naïve-groups containing same focal female at 4 and 12 weeks exposure to long-term social environments (r^2^ ± 95% confidence intervals). (**B**.) Consistency of behavior of the 3 individuals within a single naïve-group trial (ICC ± 95% confidence intervals). (**C**.) Visual representation of high intraclass correlation among individuals in the same naïve-group trial. Within each treatment category (Monoclonal, Clone 1, Clone 2) each row represents tracks of the three individuals in a single naïve-group trial. At each time point (Pre-exposure (Pre), 4 wk, and 12 wk) the focal fish and two naïve partners were tracked. For a given treatment, the same focal female was present at each time point, but her two social partners were different individuals across time points.

The mean shoaling distance in the naïve-groups was unaffected by the social environment experienced by the focal fish or duration of exposure (Table 2C, effect estimates for fixed effects in electronic supplementary material, table S8, Figure S4). There was a trend for different treatment groups to behave differently over time, but the interaction term did not reach significance (table 2C, electronic supplementary material, Figure S5). Cumulative aggression did not significantly predict shoaling distance (Table 2E; β=-0.003±0.047). Among-group variance was unaffected by treatment or exposure time (electronic supplementary material, Table S6), and group-level consistency was low for shoaling behavior (r=0.142; electronic supplementary material, Table S7).

### (d) Individuals within naïve groups behave very similarly

Focal and stimulus fish within the naïve groups were unfamiliar with one another and had different social experiences prior to the trials. Focal fish were drawn from the long-term social environments, whereas stimulus fish were all genetically identical, all of similar age and size, and all had similar prior social experience that differed substantially from that of the focal fish. Moreover, there was substantial variation in behavior across different trials, as indicated by the significant effects of long-term social environment described above. Nevertheless, the three individuals in a given naïve-group trial behaved in a remarkably similar manner. Figure 3C shows representative tracking data for 3 different trios from the naïve-group tests (see electronic supplementary material, Figure S6 for 12 additional representations). The striking visual similarity of tracking patterns within a given trial is reflected in the high ICC estimate for individual exploratory behavior (ICC=0.831 overall, values for each treatment group along with 95% CI and variance -component estimates used to calculate ICC reported in Figure 3B and electronic supplementary material, Table S9). That is, less than 17% of the total variation in behavior occurred among the three females within a given naïve-group trial, despite the substantial differences in behavior among trials that is evident in Figure 3 and electronic supplementary material, Figure S6. This high ICC value indicates that all three individuals within a given trial exhibited highly similar behavior, despite their different prior experience.

## Discussion

Elucidating the heritable causes of individual and group-emergent phenotypes is necessary to understand the evolution of social traits and other interacting phenotypes. Here, we demonstrate that phenotypic effects of genetically different social environments (IGE) carry over to a novel social environment to influence the behavior of individuals that did not experience IGE. This cascading effect is distinct from ‘second-order’ IGE [24, 25], in which the presence of genetically different individuals influences interactions between other members of the same group. Our results expand the scope of IGE by demonstrating that they can influence phenotypes even when there is no genetically-based variation present within groups. Given the prevalence of dispersal and fission-fusion group structure, there is substantial opportunity for cascading IGE in nature.

The cascading IGE we observed was associated with different levels of aggression that focal fish experienced in the long-term social environments. Somewhat surprisingly, it was the social environment containing fish of the same clone as the focal animals that exhibited the most aggression (and the naïve groups containing these focal fish exhibited the most stress behaviors). Previous studies found that Amazon mollies exhibited less aggression towards sister clones when compared to non-sister clones [43, 57]. However, a different focal clonal lineage was used in those studies, suggesting that responses to sister and non-sister clones (and, therefore, first-order and cascading IGE) vary across genotypes. Length of initial exposure may also play an important role, as in the former two studies, exposure time was short (only 10 mins or so); whereas another study using 30-days of exposure showed that familiarity influenced aggression within a clonal lineage [42]. Thus, the monoclonal environment fish may have found their naïve groups more ‘familiar’ and therefore exhibited more aggression. Furthermore, this kind of interaction between the direct effect of an individual’s genotype and IGE can produce frequency-dependent and other forms of balancing selection that can maintain, or rapidly erode genetic variation [19, 58]. The possibility that similar effects could arise from the interaction of direct genetic variance and cascading IGE warrants future empirical and theoretical investigation.

In this experiment, it is possible that the cascade of IGE that we observed occurred because focal females in Clone 1 and Clone 2 treatments experienced a genetic change in the social environment when they moved into the naïve-groups, but focal fish from the Monoclonal treatment did not. If this were the primary cause of cascading IGE, we would expect a significant difference in cascading effects between the Monoclonal treatment and both Clone 1 and Clone 2 (which we do find), but not between Clone 1 and Clone 2 (for which we found only a non-significant trend). Clone 1 and Clone 2 treatment did significantly differ in variance among naïve groups in the two treatments and in consistency of naïve-group behavior at 4 and 12 weeks. These differences in variance and consistency of behavior of naïve groups with focal females from Clone 1 versus Clone 2 suggest that sister-clone recognition was not the only cause of cascading IGE in this experiment, and points to differences in phenotypic variance as an under-explored consequence of IGE in general. In any case, our data support the hypothesis that genetically identical fish (the naïve partners) behave differently depending on genetic variation in the prior social environment experienced by another member of the group (the focal female). Whether cascading IGE depend on the degree of genetic similarity between past and current social partners should be a focus of future research. We also note that movement among groups that differ in genetic similarity to the migrating individual is likely to occur in species such as Amazon mollies that exhibit spatial population-genetic structure [35].

We detected no effects of exposure time within the long-term social environments on aggression in those environments or on cascading IGE in the naïve groups. Time-course effects on first-order IGE have been found in a related poeciliid, the eastern mosquitofish [18, 19], and increased exposure time led to higher aggression in previous studies of Amazon mollies [42, 48]. However, the time course effects of IGE reported in mosquitofish occurred during maturation, whereas the fish in our experiment were fully mature at the start of the study. The two studies that reported exposure-time effects on aggression in Amazon mollies maintained the animals at considerably higher density than that used in our experiment (48: 1.9 L / fish; 42: 4 L / fish; the present study: 6.3 L / fish), suggesting that exposure-time effects could be density-dependent.

The relatively low density in our long-term social environments might also account for lack of treatment or cascading effects on shoaling distance, despite strong effects on exploratory behavior. Anderson et al. [25] found that second-order IGE influenced social cohesion in *D. melanogaster*, and the extensive literature on leadership in social organisms indicates that differences among individual group members can substantially influence group-emergent behaviors such as shoaling [61, reviewed in 1]. We therefore expected that cascading IGE would be an important source of individual variation that generates group-emergent phenotypes [62, 63]. In our experiment, groups consisted of only 3 individuals in a small enclosure, which might limit the tendency of these fish to shoal. Experiments that use larger groups and enclosures that allow more flexibility in fission-fusion dynamics should be deployed to determine the extent to which cascading IGE influence group-emergent phenotypes.

In summary, we found that IGE propagate beyond individuals that directly experience them in Amazon mollies and possibly in many group-living species. These cascading IGE are a potentially important cause of individual differences that can lead to the emergence of leaders and followers, shoaling, swarming, and other group-emergent phenotypes. Theoretical and empirical expansion of the quantitative genetic framework developed for IGE to include cascading or other types of carry-over effects will facilitate understanding of phenotypic variation and its evolution.

## Acknowledgements

We would like to thank Mitch Daniel, Kevin Dixon, and Alexa Guerrera for their input on a previous version of this manuscript, Ryan Kelly, Hannah Lange, and Jacob Gottlieb for assistance with the behavioral setup and fish maintenance, and Christopher Schutz for assistance with the video tracking. This research was approved by the Florida State University’s Institutional Animal Care and Use Committee (#1704 and 201900038). AMM was supported by Provost Postdoctoral Fellowship Program, DB was funded by the German Research Foundation (DFG) through Germany’s Excellence Strategy (EXC 2002/1 ‘Science of Intelligence’, project number 390523135), and KH was funded by NSF IOS 1354775 and NSF DEB 1740466.

## Notes

### Competing Interest Statement

The authors have declared no competing interest.

