## Supplemental Methods for "Cascading indirect genetic effects in a clonal vertebrate"

### 1. Material and methods

*(a) Study Specimens*

Two weeks prior to initiating the experiment, we marked all fish with elastomer tags (two 3mm subcutaneous marks anterior and/or posterior of the dorsal fin) to allow us to identify focal individuals from partner females. To recover, fish were placed in 113.6L aquaria treated with Stress Coat+® (API®) and sea salt (Instant Ocean®), with each aquarium containing an average of ten sister clone individuals. Females were all virgins and, thus, were all receptive but not pregnant at the time of the trials. They were fed daily ad libitum with commercial fish food (Tetramin® tropical flakes). Experiments occurred from August to November 2019, and fish were exposed to natural light cycles during the course of the experiment.

*(b) Long-term social environments*

All aquaria contained sand and two small PVC pipe fittings (2 cm) for shelter, with one long side and two short sides covered with blue tarp to prevent visual communication with neighboring tanks.

We excluded one datum (block 7, clone 1 treatment, week 3) because it was an extreme outlier: 174 aggressive acts (2.3x higher than the maximum number of acts in any other tank), and 190 s of total aggression (2.2x larger than the maximum time spent being aggressive in any other tank). Individual identification was not possible during the trial while fish were in motion and visible only from one side. Therefore, we used the total number and duration of these behaviors across all fish in the trial as a measure of the social environment within the tank.

*(c) Naïve-group tests*

This test chamber was placed inside a frame covered with blue tarp to minimize external disturbance. All water in the chamber was replaced with clean freshwater prior to every test. All videos were edited to remove the first and last 2 minutes of recording (VideoPad Video Editor by NCH sofware©, v. 8.40) to allow for acclimation to the experimental tank and to remove any influence of camera or experimenter movement at the beginning and end of trial (i.e., leaving or approaching the experimental setup to turn on/off camera). All cropped videos were 6 minutes long and were analyzed by a blind observer using EthoVision XT (Noldus, v14). Within the EthoVision program, we distinguished the three individuals throughout the analyses and acquired movement and position data (Cartesian coordinates) for all three individuals. Although fish could be individually tracked, the focal individual could not be distinguished from the novel partner fish on the videos; therefore, we did not calculate separate metrics for focal and novel partner fish. We extracted the following measures from EthoVision: distance traveled (cm), velocity ($cm/s^2$), frequency entering white zone, duration in white zone (s), latency to enter white zone (s), frequency entering grey zone, duration in grey zone (s), time spent immobile (s; freezing behavior), and distance between individuals (cm; shoaling distance).

*(d) Ethics*

All fish tanks included substrate and enrichment and were maintained with weekly water changes throughout the duration of this study. Fish never suffered from food deprivation or injuries during this study.

*(e) Analyses*

*(e.1) Long-term social environment groups.*

Initial models included fixed effects of exposure time (weeks), treatment-by-time interaction, the baseline (week 0) measure of aggression PC1, focal female standard length (log-transformed), and the average standard length of the social-partner females (log-transformed). A random group ID effect was used to account for repeated measures on groups (at weeks 4 and 12); initial models also included a random effect due to experimental block. Baseline aggression and size of social partners never approached significance in initial models (electronic supplemental material, Table S3A), and the random block effect was consistently near zero and never significant. These terms were therefore not included in the final models. Random-effect estimates (variance among groups at each time point and the covariance of measures of the same group across time points) were allowed to vary by treatment group in initial models (using the *group* option). If allowing variance to be group specific did not significantly improve the fit of the model (by a likelihood ratio test), the final model assumed that treatment groups shared the same variance-covariance structure.

*(e.2) Naïve-group tests.*

Frequency entering and duration in the gray zone was redundant with information for entering and duration in the white zone, so we used only the data for the white zone in the analyses.

*Average behavior of naïve-groups.* T

Initial models included fixed effects of the long-term social environment of the focal fish, time in the long-term environment, size of the focal female, the mean size of the novel partner females, mean size of the long-term social partners (all size variables were log-transformed), and baseline behavior (at week 0). To assess whether the effect of the long-term social environment was mediated by aggression in that environment, initial models also included a measure of the sum of all aggression experienced up to the time the assay was conducted (e.g., either 0 to 4 weeks or 0 to 12 weeks). Initial models also included a random effect due to experimental block. The block random effect was always near zero and never approached significance, so it was dropped from the final models.
