## Supplemental tables and figures for "Cascading indirect genetic effects in a clonal vertebrate"

**Supplemental Table S1:** Test statistic and p-value for statistical model of log-transformed focal female standard length, mean social partner female standard length, and mean naïve-group partner standard length as dependent variables.

| Model | Effect | Statistic | P-value |
|---|---|---|---|
| ***A. Focal female standard length*** | | | |
| | Social environment | $F_{2,33} = 0.230$ | 0.7986 |
| ***B. Mean social partner female standard length*** | | | |
| | Social environment | $F_{2,33} = 2.56$ | 0.0927 |
| ***C. Mean naïve-group partner standard length*** | | | |
| | Social environment | $F_{2,33} = 1.86$ | 0.1709 |

**Supplemental Table S2:** Least-squares mean ± SE for focal female standard length, mean social partner female standard length, and mean naïve-group partner standard length across each treatment.

| Measurement | Treatment | Average SL (mm) | LS mean ± SE | Test effect |
|---|---|---|---|---|
| ***A. Focal female standard length*** | | | | $F_{2,33} = 0.23$, $p = 0.7986$ |
| | Monoclonal | 30.83 ± 2.92 | 1.5094 ± 0.0379 | |
| | Clone 1 | 32.94 ± 1.85 | 1.5438 ± 0.0379 | |
| | Clone 2 | 32.39 ± 2.06 | 1.5169 ± 0.0379 | |
| ***B. Mean social partner female standard length*** | | | | $F_{2,33} = 2.56$, $p = 0.0927$ |
| | Monoclonal | 32.17 ± 2.15 | 1.5226 ± 0.0076 | |
| | Clone 1 | 33.96 ± 1.71 | 1.5065 ± 0.0076 | |
| | Clone 2 | 33.38 ± 2.08 | 1.5304 ± 0.0076 | |
| ***C. Mean naïve-group partner standard length*** | | | | $F_{2,33} = 1.86$, $p = 0.1709$ |
| | Monoclonal | 31.32 ± 3.02 | 1.5118 ± 0.0086 | |
| | Clone 1 | 32.90 ± 2.43 | 1.4938 ± 0.0086 | |
| | Clone 2 | 32.61 ± 2.72 | 1.5160 ± 0.0086 | |

**Supplemental Table S3:** Test statistic and p-value for full statistical model of aggression PC1, exploratory/stress behaviors, and shoaling behaviors as dependent variables.

| Model | Effect | Statistic | P-value |
|---|---|---|---|
| ***A. Aggressive behavior in long-term social treatments (PC1)*** | | | |
| | Baseline aggression | $F_{1,197.4} = 0.05$ | 0.824 |
| | **Focal female standard length** | $\mathbf{F_{1,194.1} = 4.33}$ | **0.039** |
| | Mean social partner standard length | $F_{1,182.6} = 0.09$ | 0.345 |
| | **Social environment** | $\mathbf{F_{2,156.9} = 6.30}$ | **0.002** |
| | Exposure time | $F_{7,218.5} = 1.29$ | 0.258 |
| | Social environment*Time | $F_{14,189.6} = 0.69$ | 0.786 |
| ***B. Exploratory behavior in naÏve-group trials (PC1)*** | | | |
| | Baseline exploration | $F_{1,18.38} = 0.00$ | 0.952 |
| | Focal female standard length | $F_{1,18.89} = 2.98$ | 0.101 |
| | Mean naïve-group partner standard length | $F_{1,33.75} = 3.37$ | 0.075 |
| | Mean social partner standard length | $F_{1,23.65} = 1.59$ | 0.219 |
| | PC1 aggression experienced | $F_{1,13.11} = 0.34$ | 0.572 |
| | **Social environment** | $\mathbf{F_{2,17.97} = 3.94}$ | **0.038** |
| | Exposure time | $F_{1,22.65} = 1.03$ | 0.320 |
| | Social environment*Time | $F_{2,17.89} = 0.64$ | 0.539 |
| ***C. Shoaling distance in naÏve-group trials*** | | | |
| | Baseline aggression | $F_{1,23.11} = 0.91$ | 0.349 |
| | Focal female standard length | $F_{1,33.18} = 0.19$ | 0.666 |
| | Mean naïve-group partner standard length | $F_{1,42.59} = 0.23$ | 0.633 |
| | Mean social partner standard length | $F_{1,19.16} = 1.10$ | 0.308 |
| | PC1 aggression experienced | $F_{1,32.49} = 0.26$ | 0.617 |
| | Social environment | $F_{2,20.46} = 0.99$ | 0.390 |
| | Exposure time | $F_{1,42.1} = 0.00$ | 0.978 |
| | Social environment*Time | $F_{2,20.75} = 2.89$ | 0.078 |

**Supplemental Table S4:** Fixed effect estimates from final GLMM on aggression PC1.

| Effect | Treatment | Week | Estimate | Error | DF | t-value | P-value |
|---|---|---|---|---|---|---|---|
| Intercept | | | -6.248 | 3.730 | 209.6 | -1.67 | 0.095 |
| Focal female SL | | | 1.931 | 1.069 | 208.1 | 1.81 | 0.072 |
| Social environment | Monoclonal | | -0.461 | 0.566 | 246.7 | -0.81 | 0.416 |
| | Clone 1 | | -0.970 | 0.565 | 246.5 | -1.72 | 0.088 |
| | Clone 2 | | 0 | - | - | - | - |
| Week | 1 | | -0.157 | 0.564 | 246.1 | -0.28 | 0.781 |
| | 2 | | -0.236 | 0.564 | 246.1 | -0.42 | 0.676 |
| | 3 | | -0.314 | 0.564 | 246.1 | -0.56 | 0.578 |
| | 4 | | -0.949 | 0.564 | 246.2 | -1.68 | 0.094 |
| | 6 | | 0.299 | 0.564 | 247.1 | -0.53 | 0.597 |
| | 8 | | 0.721 | 0.564 | 254.2 | -1.28 | 0.202 |
| | 10 | | -0.944 | 0.545 | 236.1 | -1.73 | 0.085 |
| | 12 | | 0 | - | - | - | - |
| Social environment*Week | Monoclonal | 1 | 0.685 | 0.798 | 246.1 | 0.86 | 0.392 |
| | Monoclonal | 2 | 0.889 | 0.798 | 246.1 | 1.11 | 0.266 |
| | Monoclonal | 3 | 1.026 | 0.798 | 246.1 | 1.29 | 0.200 |
| | Monoclonal | 4 | 1.141 | 0.798 | 246.2 | 1.43 | 0.154 |
| | Monoclonal | 6 | 0.921 | 0.798 | 247.1 | 1.15 | 0.249 |
| | Monoclonal | 8 | 1.013 | 0.797 | 254.2 | 1.27 | 0.205 |
| | Monoclonal | 10 | 1.354 | 0.771 | 236.1 | 1.76 | 0.080 |
| | Monoclonal | 12 | 0 | - | - | - | - |
| | Clone 1 | 1 | 0.998 | 0.798 | 246.1 | 1.25 | 0.212 |
| | Clone 1 | 2 | 0.196 | 0.798 | 246.1 | 0.25 | 0.806 |
| | Clone 1 | 3 | 1.110 | 0.806 | 248.5 | 1.38 | 0.170 |
| | Clone 1 | 4 | 0.916 | 0.798 | 246.2 | 1.15 | 0.252 |
| | Clone 1 | 6 | -0.143 | 0.798 | 247.1 | -0.18 | 0.858 |
| | Clone 1 | 8 | 0.981 | 0.797 | 254.2 | 1.23 | 0.219 |
| | Clone 1 | 10 | 0.566 | 0.771 | 236.1 | 0.73 | 0.463 |
| | Clone 1 | 12 | 0 | - | - | - | - |
| | Clone 2 | 1 | 0 | - | - | - | - |
| | Clone 2 | 2 | 0 | - | - | - | - |
| | Clone 2 | 3 | 0 | - | - | - | - |
| | Clone 2 | 4 | 0 | - | - | - | - |
| | Clone 2 | 6 | 0 | - | - | - | - |
| | Clone 2 | 8 | 0 | - | - | - | - |
| | Clone 2 | 10 | 0 | - | - | - | - |
| | Clone 2 | 12 | 0 | - | - | - | - |

**Supplemental Table S5:** Fixed effect estimates from GLMM on exploratory/stress behaviors PC1 in the naïve-group trials.

| Effect | Treatment | Week | Estimate | Error | DF | t-value | P-value |
|---|---|---|---|---|---|---|---|
| Intercept | | | 0.241 | 0.242 | 20.63 | 1.00 | 0.330 |
| Social environment | Monoclonal | | -0.978 | 0.312 | 36.01 | -3.13 | 0.003 |
| | Clone 1 | | -0.248 | 0.377 | 30.55 | -0.66 | 0.516 |
| | Clone 2 | | 0 | - | - | - | - |
| Week | 4 | | 0.143 | 0.295 | 11.00 | 0.48 | 0.638 |
| | 12 | | 0 | - | - | - | - |
| Social environment*Week | Monoclonal | 4 | 0.319 | 0.338 | 17.31 | 0.94 | 0.359 |
| | Monoclonal | 12 | 0 | - | - | - | - |
| | Clone 1 | 4 | -0.076 | 0.616 | 17.01 | -0.12 | 0.903 |
| | Clone 1 | 12 | 0 | - | - | - | - |
| | Clone 2 | 4 | 0 | - | - | - | - |
| | Clone 2 | 12 | 0 | - | - | - | - |

**Supplemental Table S6:** Tests of covariance parameters based on the restricted likelihood ratio for exploratory behavior but not for shoaling behavior.

| **Model** | | **-2 Res Log Like** | **$X^2$ value** | **DF** | **Significance** |
|---|---|---|---|---|---|
| *Exploratory/stress behavior in naïve-group trials (PC1)* | Variance differs by group? | 180.04 | 26.00 | 6 | p = 0.0002 |
| | Repeatability significant? | 172.79 | 18.76 | 3 | p = 0.0003 |
| | Complete lack of variance structure? | 172.79 | 18.76 | 3 | p = 0.0003 |
| *Shoaling distance in naïve-group trials* | Variance differs by group? | -46.30 | 6.28 | 4 | p = 0.179 |
| | Repeatability significant? | -51.88 | 0.69 | 3 | p = 0.875 |
| | Complete lack of variance structure? | -51.88 | 0.69 | 3 | p = 0.875 |

**Supplemental Table S7:** Consistency of behavior for naïve groups across 4 and 12 weeks measured using Pearson's r and 95% confidence limits. Variance differed by treatment for exploratory behavior but not for shoaling behavior.

| Behavior | Social environment | Pearson's r | Lower 95% CL | Upper 95% CL | Significance |
|---|---|---|---|---|---|
| PC1 exploratory | Monoclonal | 0.732 | 0.272 | 0.920 | Z = 5.23, p < 0.0001 |
| | Clone 1 | -0.759 | -0.887 | -0.023 | Z = -5.92, p < 0.0001 |
| | Clone 2 | 0.280 | -0.350 | 0.736 | Z = 1.01, p = 0.314 |
| Shoaling | Monoclonal | 0.190 | -0.427 | 0.691 | Z = -0.02, p = 0.985 |
| | Clone 1 | 0.176 | -0.443 | 0.682 | Z = 0.62, p = 0.533 |
| | Clone 2 | -0.006 | -0.578 | 0.570 | Z = 0.52, p = 0.604 |

**Supplemental Table S8:** Fixed effect estimates from GLMM on shoaling behavior in the naïve-group trials.

| Effect | Treatment | Week | Estimate | Error | DF | t-value | P-value |
|---|---|---|---|---|---|---|---|
| Intercept | | | 0.808 | 0.037 | 21.46 | 22.05 | <0.0001 |
| Social environment | Monoclonal | | 0.080 | 0.051 | 43.42 | 1.42 | 0.128 |
| | Clone 1 | | -0.034 | 0.068 | 36.39 | -0.51 | 0.613 |
| | Clone 2 | | 0 | - | - | - | - |
| Week | 4 | | 0.063 | 0.048 | 11.00 | 1.33 | 0.212 |
| | 12 | | 0 | - | - | - | - |
| Social environment*Week | Monoclonal | 4 | -0.164 | 0.070 | 21.89 | -2.35 | 0.028 |
| | Monoclonal | 12 | 0 | - | - | - | - |
| | Clone 1 | 4 | 0.002 | 0.086 | 19.05 | 0.02 | 0.981 |
| | Clone 1 | 12 | 0 | - | - | - | - |
| | Clone 2 | 4 | 0 | - | - | - | - |
| | Clone 2 | 12 | 0 | - | - | - | - |

**Supplemental Table S9:** Intraclass correlation coefficient and variance components for PC1 for exploratory behaviors, along with corresponding residuals and their tests statistics. There was no significant difference among the three social environment treatments.

| Model | Social environment | Parameter | Variance | Significance | ICC | Lower 95% CI | Upper 95% CI |
|---|---|---|---|---|---|---|---|
| PC1 for exploratory behavior | Overall | Group | 0.836 ± 0.122 | Z = 6.84, p < 0.0001 | 0.831 | 0.781 | 0.882 |
| | | Residual | 0.170 ± 0.016 | Z = 10.39, p < 0.0001 | | | |
| | Monoclonal | Group | 0.741 ± 0.186 | Z = 3.99, p < 0.0001 | 0.829 | 0.738 | 0.920 |
| | | Residual | 0.131 ± 0.022 | Z = 6.00, p < 0.0001 | | | |
| | Clone 1 | Group | 0.973 ± 0.246 | Z = 3.95, p < 0.0001 | 0.825 | 0.730 | 0.919 |
| | | Residual | 0.205 ± 0.034 | Z = 6.00, p < 0.0001 | | | |
| | Clone 2 | Group | 0.793 ± 0.201 | Z = 3.95, p < 0.0001 | 0.852 | 0.767 | 0.936 |
| | | Residual | 0.172 ± 0.029 | Z = 6.00, p < 0.0001 | | | |

**Supplemental Figure S1:** Heatmap of Pearson's correlations among exploratory behavioral variables at the group level (**A.**) and at the individual level (**B.**).

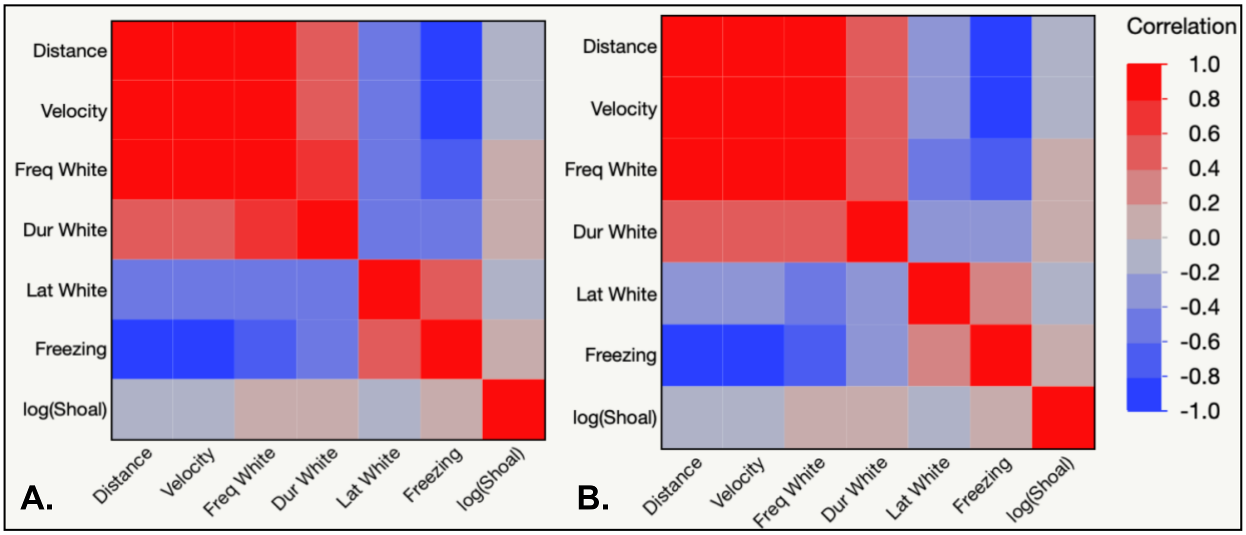

**Supplemental Figure S2:** Average (white triangles) ± standard error for the PC1 score summarizing the amount of aggression experienced within each social environment. (Monoclonal – purple; Clone 1 – teal; Clone 2 – yellow). Violin plots depict the distribution of the PC1 scores for each treatment after the first four weeks and after 12 weeks of exposure, with raw scores represented by each open circle.

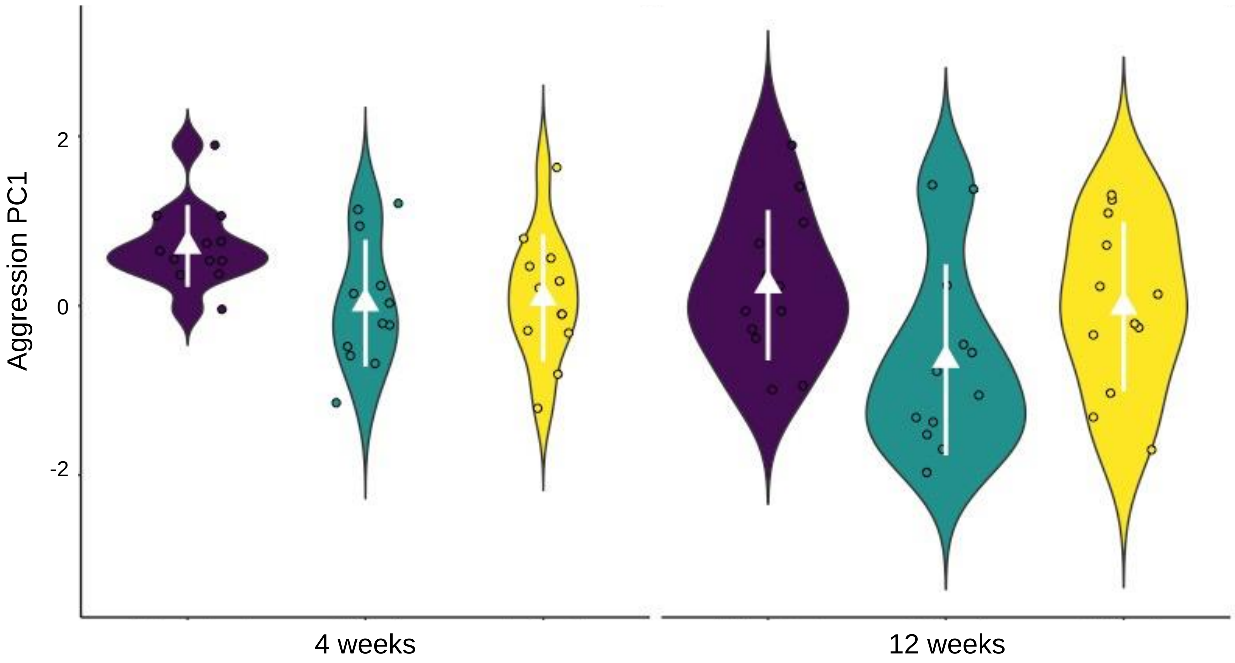

**Supplemental Figure S3:** Average (white triangles) ± standard error for the PC1 score on the exploratory behaviors within each social environment. (Monoclonal – purple; Clone 1 – teal; Clone 2 – yellow). Violin plots depict the distribution of the PC1 scores for each treatment after the first four weeks and after 12 weeks of exposure, with raw scores represented by each open circle.

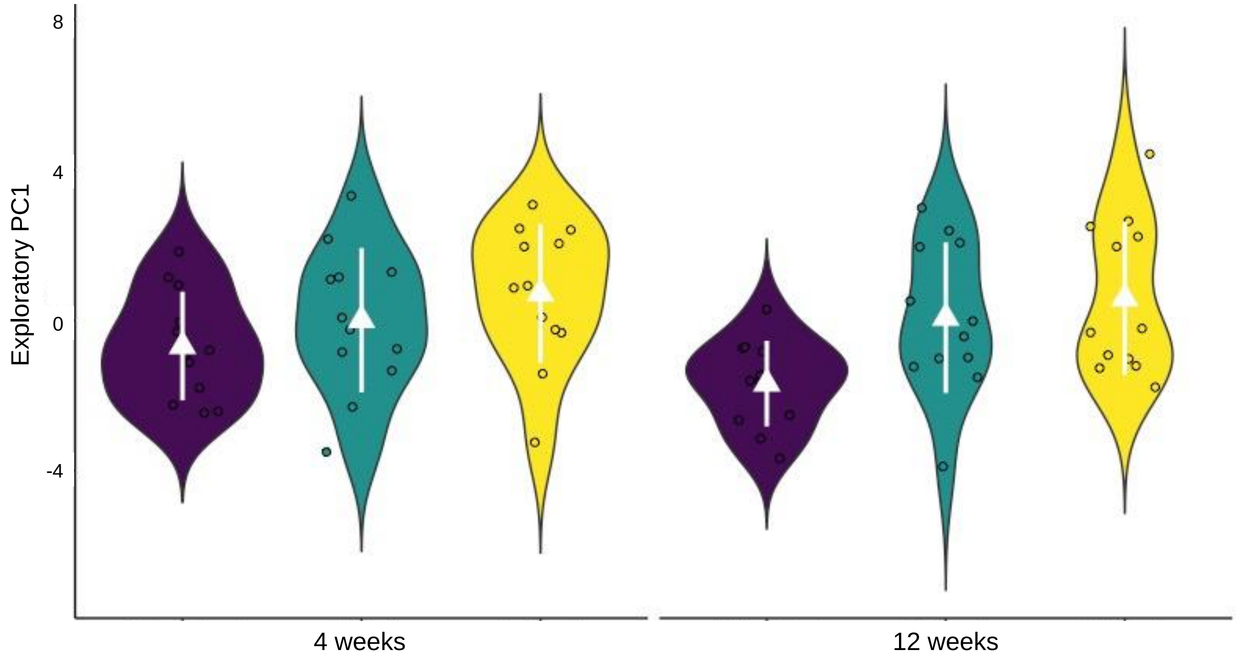

**Supplemental Figure S4:** Average (white triangles) ± standard error for shoaling behavior within each social environment. (Monoclonal – purple; Clone 1 – teal; Clone 2 – yellow). Violin plots depict the distribution of shoaling for each treatment after the first four weeks and after 12 weeks of exposure, with raw scores represented by each open circle.

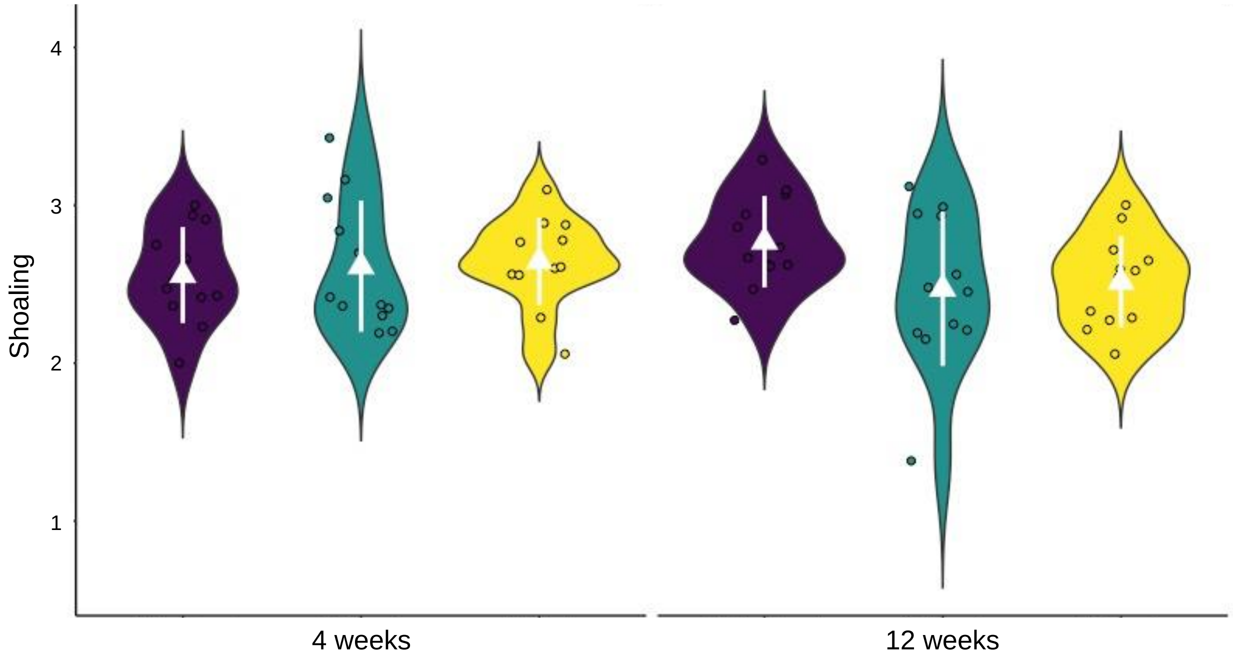

**Supplemental Figure S5:** Least square means +/- standard error from ln(shoaling behavior) for the interaction between the long-term social environment a focal female experienced and the duration of exposure to that environment (squares and dotted line = 4 weeks; circles and dashed line = 12 weeks) (Monoclonal = blue; Clone 1 = yellow; Clone 2 = pink).

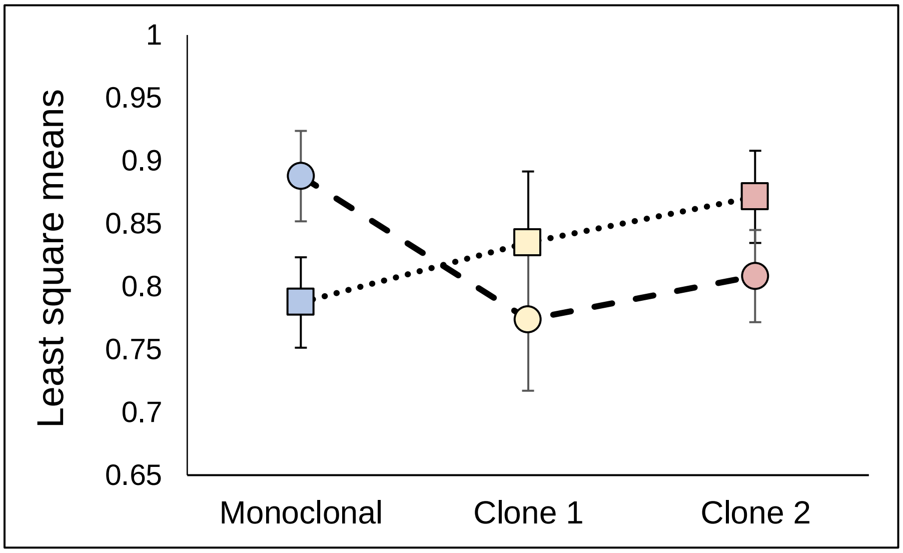

**Supplemental Figure S6:** Representative heatmaps of four focal females from each of the different social environments. Within each treatment are the heatmaps of the three individuals (i.e., the focal female + 2 naïve sister clones; S1-S36) within each naïve-group trial before exposure to the social environment (Pre), 4 weeks (4 wks), and 12 weeks (12 wks) after exposure to each social environment (monoclonal, clone 1, and clone 2).

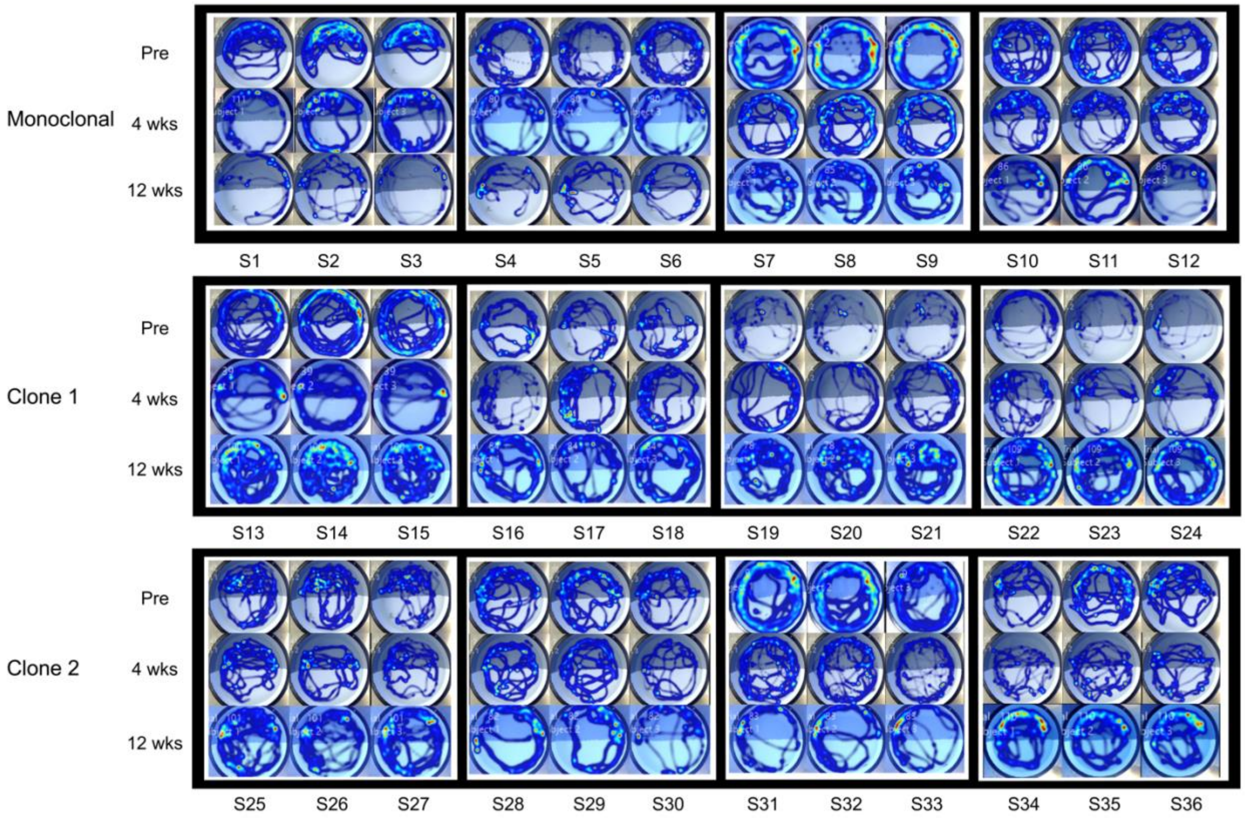
